# Superior Colliculus-Projecting GABAergic Retinal Ganglion Cells Mediate Looming-Evoked Flight Response

**DOI:** 10.1101/2020.05.14.090167

**Authors:** Xue Luo, Danrui Cai, Kejiong Shen, Qinqin Deng, Xinlan Lei, Sen Jin, Wen-Bo Zeng, Hua Li, Fuqiang Xu, Lu Huang, Chaoran Ren, Min-Hua Luo, Ting Xie, Yin Shen

## Abstract

The looming stimulus-evoked flight response is an experimental paradigm for studying innate defensive behaviors. However, how the visual looming stimulus is transmitted from the retina to the brain remains poorly understood. Here, we report that superior colliculus (SC)-projecting RGCs transmit the looming signal from the retina to the brain to mediate the looming-evoked flight behavior by releasing GABA. In the mouse retina, GABAergic RGCs are capable of projecting to many brain areas, including the SC. Superior colliculus (SC)-projecting GABAergic RGCs (spgRGCs) are mono-synaptically connected to the parvalbumin-positive SC neurons known to be required for the looming-evoked flight response. Optogenetic activation of spgRGCs triggers GABA-mediated inhibition in SC neurons. The ablation or silence of spgRGCs compromises looming-evoked flight response but not image-forming functions. Therefore, this study shows that spgRGCs control the looming-evoked flight response by regulating SC neurons via GABA, providing novel insight into the regulation of innate defensive behaviors.

## Introduction

The ability to detect the naturally occurring looming stimulus, which mimics approaching objects, is a conserved innate defensive behavior important for the survival of animal species, ranging from *Drosophila* to mammals, including humans [1–3]. The looming stimulus-induced flight response in mice has been recently established as an effective model for studying innate defensive behaviors in animals [4–6]. The neural circuits for mediating the looming-evoked innate defensive behavior in the mouse brain have recently been shown to involve different brain areas, including the superior colliculus (SC). However, how the looming stimulus signal is transmitted from the retina to the brain remains largely elusive.

The looming-evoked innate defensive behavior has recently been shown to involve the brain neural circuits mediated through the SC, the parabigeminal nucleus (PBGN), lateral posterior thalamic nucleus (LPTN), the ventral tegmental area (VTA), the zona incerta (ZI) and the amygdala in the mouse [4]. The SC is generally believed to serve as a sensory-motor center for processing and controlling innate visual behaviors [7, 8]. Parvalbumin positive (PV^+^) glutamatergic excitatory neurons in the superficial layer of the SC project to different brain areas to mediate the looming-evoked escape and freezing behavior [4]. Interestingly, the looming stimulus simultaneously activate two distinct groups of PV^+^ GABAergic SC neurons projecting the PBGN and the LPTN, which both innervate the amygdala where the fear response is initiated, to mediate escape and freezing responses, respectively [9]. Also, glutamatergic SC neurons innervate and send excitatory signals to GABAergic neurons in the VTA, which further project to the central nucleus of the amygdala, mediating the looming-evoked defensive behavior [10]. Although GABAergic neurons in the ZI directly innervate excitatory neurons in the dorsolateral and ventrolateral compartments of periaqueductal gray (PAG) to drive escape and freezing responses, it remains unclear how these neurons are connected to the SC [11].

The visual looming stimulus must be transmitted through retinal ganglion cells (RGCs) since they are the only cell type connecting the eye to the brain. Interestingly, GABAergic dorsal raphe nuclei (DRN) neurons antagonize the serotonergic DRN neurons to activate the SC-LPTN-amygdala pathway to mediate the looming-evoked flight response [6]. It still remains unclear how the SC/DRN-projecting RGCs transmit the looming stimulus signal. Recently a subset of melanopsin-expressing intrinsically photosensitive RGCs (ipRGCs) in mice that release GABA at the non-image-forming brain targets, including suprachiasmatic nucleus (SCN) and olivary pretectal nucleus (OPN), was identified. Meanwhile this GABAergic circuit decreases the sensitivity of the non–image-forming visual system at low light levels [12]. Here we identified SC-projecting GABAergic RGCs (spgRGCs) transmit the looming stimulus signal and mediate the looming-evoked flight response by releasing GABA.

## Results

### The mouse retina contains a population of GABAergic RGCs projecting to different regions of the brain

Earlier studies suggest the existence of GABA-immunoreactive RGCs in turtle [13], rabbit [14, 15] and . and mouse retinas. To furthermore confirm the existence and functions of GABAergic RGCs in the mouse retina, the recombinant adeno-associated virus (AAV) expressing EYFP-EYFP in a Cre-dependent manner were injected into the vitreous of adult eyes of *Gad2-Cre* or *vGAT-Cre* mice, which are specifically expressed in GABAergic neurons [16–20] (Fig 1A). In addition to EYFP-labeled GABAergic amacrine cells in the inner nuclear layer (INL), EYFP-labeled GABAergic RGCs in the ganglion cell layer (GCL) also existed based on their EYFP-positive optic nerve and laminar positions [21–23] (Fig 1B). Those EYFP-labeled RGCs were not caused by virus leakage since there were no EYFP-labeled retinal cells in negative controls (S1 Fig). Consistently, 13% of RGCs (n=6225) are positive for *Gad2* mRNA expression based on the previously published scRNA results [24] (Fig 1C). Those labeled YFP-labeled GABAergic RGCs project their axons in the optic track (Opt) in the brain, and mainly innervate the SCN, the lateral geniculate nucleus (LGN) and the SC, but not the dorsal raphe nucleus (DRN) (Fig 1D-H). In the LGN, the axonal arbors of the GABAergic RGCs spread across the LGN, but appear to be more abundant in the dorsal side (DLG) and intergeniculate leaflet (ILG) than in the ventral side (VLG) (Fig 1F). In the SC, the axonal arbors of GABAergic RGCs are more restricted to the superficial and intermediate layers (SL and IL) in the SC (Fig 1G). Taken together, GABAergic RGCs exist in the adult mouse eye and are capable of projecting to the brain areas participating in image-forming and non-image-forming functions.

**Fig 1.**
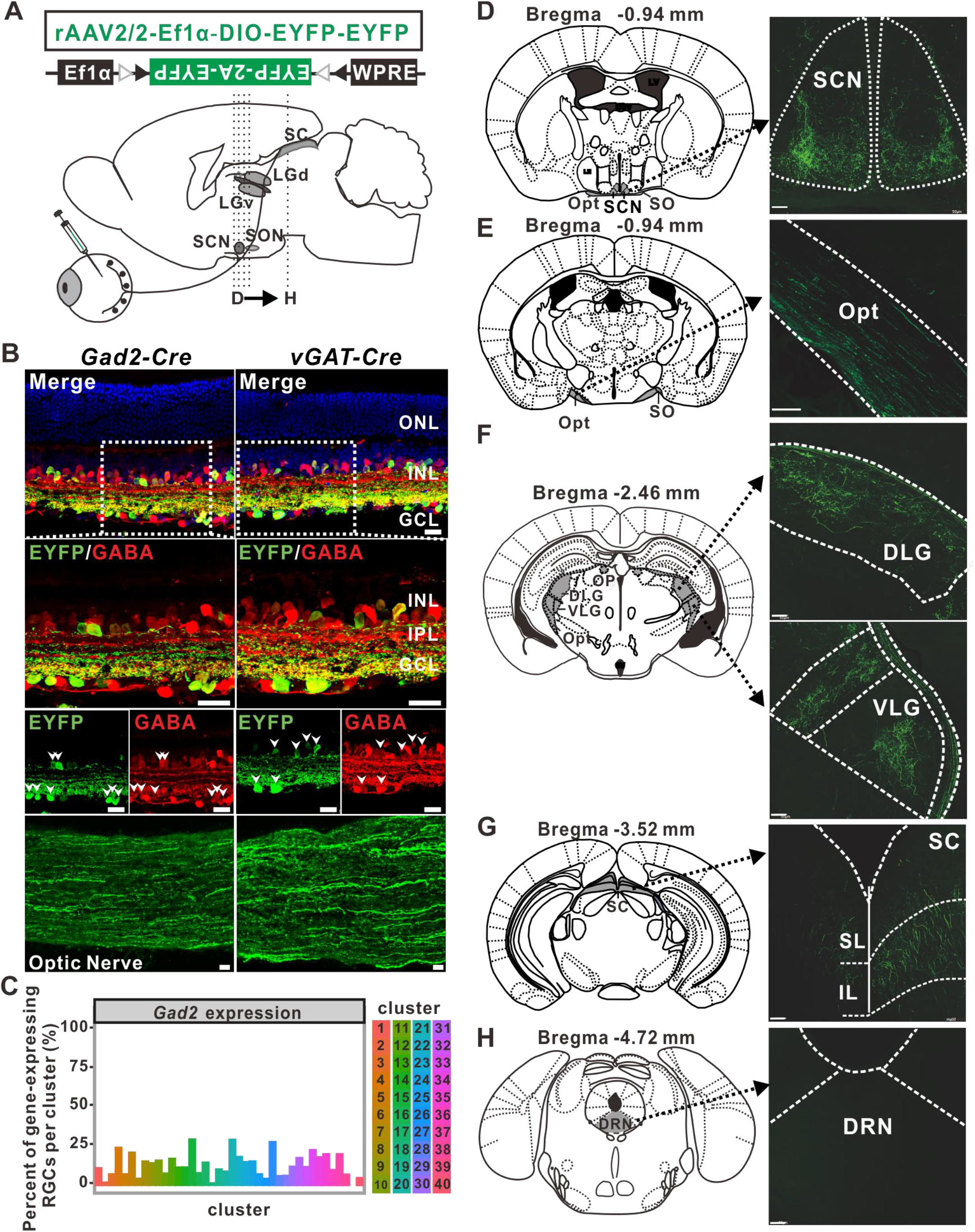
GABAergic RGCs project to multiple areas of the mouse brain. (**A**) Schematic diagrams explaining the anterograde labeling of GABAergic RGCs and the gene structure for Cre-dependent expression of two copies of EYFP in the AAV. (**B**) EYFP-labeled GABAergic neurons in the *Gad2-Cre* (left) and *vGAT-Cre* (right) retinas. EYFP-labeled RGCs are expressing gamma-aminobutyric acid (GABA) (shown in high magnifications, 100X), and can also extend EYFP-positive axons in the optic nerve (ONL: outer nuclear layer; INL: inner nuclear layer; GCL: ganglion cell layer; Scale bar: 20 μm). (**C**) *Gad2* mRNA expression in mouse RGC subpopulations based on the previously published scRNA results. (**D-H**) Brain projections from the EYFP-labeled RGCs revealed by anterograde labeling. These retinorecipient brain regions (shown in high magnifications; right) correspond to the green-filled areas in the mouse brain atlas (left) (SCN: suprachiasmatic nucleus; Opt: optic tract; SC: superior colliculus; DLG: dorsal lateral geniculate nucleus; IGL: intergeniculate leaf; VLG: ventral lateral geniculate nucleus; DRN: dorsal raphe nucleus. Scale bar: 100 μm).

### spgRGCs are capable of releasing GABA to elicit physiological responses in SC neurons

To determine if spgRGCs can functionaly release GABA to elicit post-synaptic responses in SC neurons, channel rhodopsin 2 and EYFP fusion protein (ChR2-EYFP) were expressed in the AAV-infected GABAergic RGCs. When spgRGCs were activated by 470 nm laser light, post-synaptic SC responses were recorded (Figs 2A and 2B). Brief pulses of light can induce sustained inward currents in the ChR2-EYFP-expressing RGCs, indicating that ChR2-EYFP can be functional in GABAergic RGCs (Fig 2C). In the SC, light-mediated ChR2 activation in the ChR2-EYFP-expressing RGC axonal fibers induces inhibitory postsynaptic currents (IPSCs) when V_holding_ was set to 0 mV, which is close to the reveasl potential for cations [25] (Figs 2D and 2E). The addition of bicuculline (10 μM), GABAA receptor antagonist, to the recording bath solution (TTX+D-AP5+CNQX) could reversibly suppress the light-evoked responses (n=26; 21.7±4.2 pA) (Fig 2E). To further characterize the properties of ChR2-induced currents, in all the neurons that we recorded, we also found ChR2-evoked excitatory (−70 mV) and inhibitory (0 mV) currents in some postsynaptic neurons (n=14) in the presence of TTX (Figs 2F and 2G). Our findings indicate that spgRGCs innervate and functionally inhibit SC neurons through releasing GABA.

**Fig 2.**
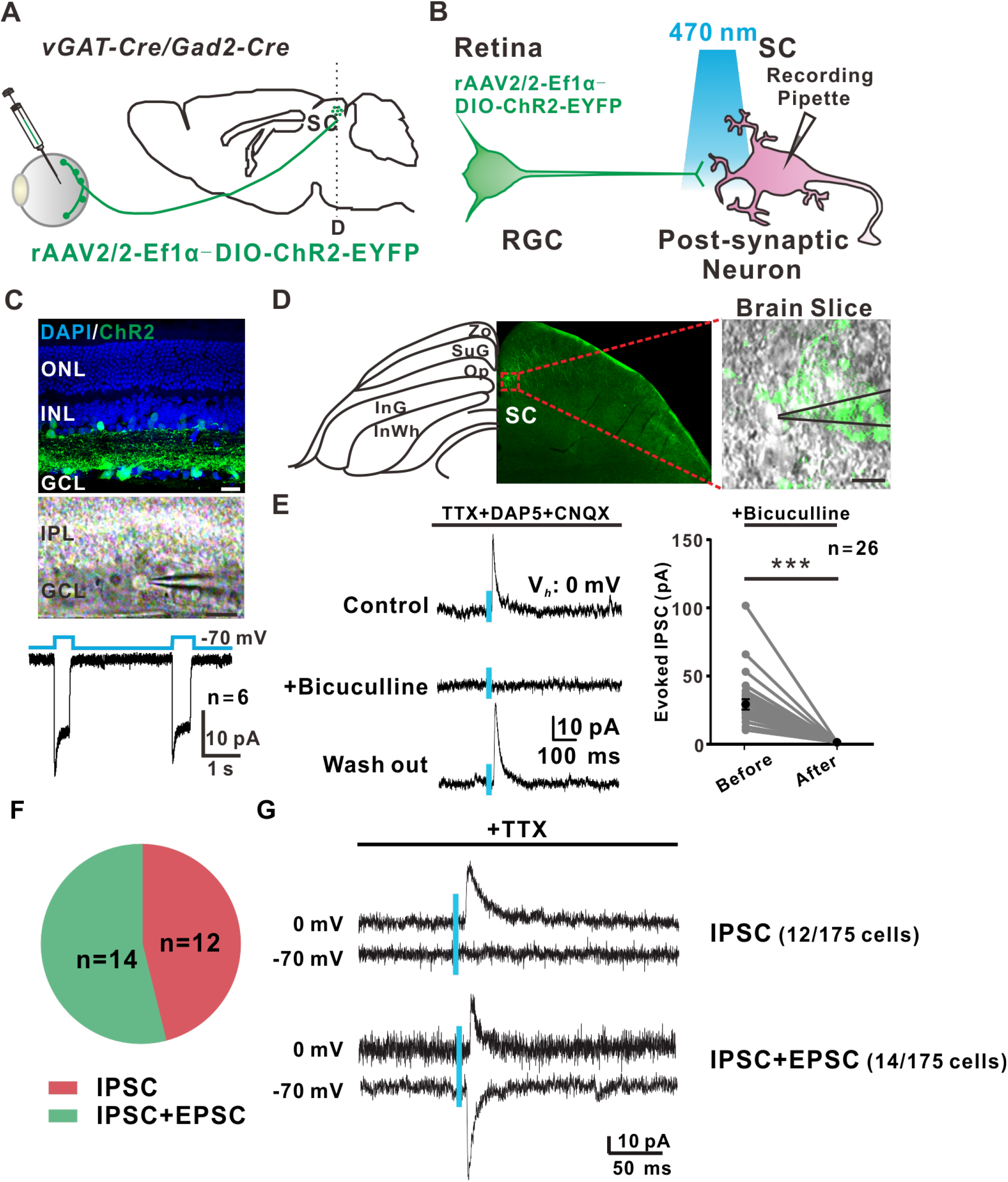
ChR2-mediated activation causes GABAergic RGCs to release GABA and evoke electrophysiological responses in SC neurons. (**A, B**) Schematic representation of optogenetic stimulation of GABAergic RGCs and recordings in labelled RGCs and SC neurons in tissue slices. (**C**) Patch-clamp recording of the AAV-ChR2-EYFP infected RGCs in the *vGAT-Cre*/*Gad2-Cre* retina. Light stimulation of ChR2 (470 nm, 500 ms) induces typical inward currents in the ChR2-EYFP-expressing RGCs. Scale bar: 25 μm. (**D**) Recording of an SC neuron receiving inputs from GABAergic RGCs (scale bar: 10 μm). (**E**) Postsynaptic IPSCs of a GABAergic RGC-projecting SC neuron in response to ChR2-mediated activation of RGCs (470nm blue light for 5 ms) are reversibly inhibited by GABAA receptor antagonist Bicuculline (10 μM). All recordings are performed with the 0 mV holding potential (V_holding_) in the presence of D-AP5, strychnine, CNQX and TTX. Right: quantification results on IPSCs. Light stimulation (470 nm) is marked by a blue bar. (F) The properties of ChR2-induced currents. IPSC/(IPSC+EPSC) ratios for ChR2^+^ neurons are analyzed from vGAT-Cre/Gad2-Cre mice. (G) Example averaged traces of ChR2-evoked IPSC (upper) and IPSC+EPSC (lower). IPSC: ChR2 evoked excitatory (−70 mV), but no inhibitory (0 mV) currents in the presence of TTX; IPSC+EPSC: ChR2-evoked excitatory (−70 mV) and inhibitory (0 mV) currents in the presence of TTX. Statistical analysis: one-way Student’s t-test; ***: *P*< 0.001.

### spgRGCs are composed of ON, OFF and ON-OFF types

To further characterize the distribution, dendritic morphology and light responses of the spgRGCs, Cre-dependent EYFP-EYFP-expressing AAV viruses were injected into the SC region of the *vGAT-Cre* or *Gad2-Cre* brain for retrogradely labeling spgRGCs (Figs 3A and 3B). Retrogradely labeled RGCs are GABAergic since the EYFP-labeled RGCs express GABA (n=12 mice) (Fig 3C). By comparing to the Brn3a-positive (Brn3a^+^) RGCs (Center: 2070.25±166.45 per mm^2^; Midperiphery: 2105.09±158.21 per mm^2^; Periphery: 1550.68± 102.62 per mm^2^), which are more dense in the central retina than in the peripheral retina [26], the EYFP-labeled spgRGCs showed a higher density in the peripheral retina (220.39±34.86 per mm^2^) than in the central retina (152.78±27.47 per mm^2^) (Figs 3D-F) and midperipheral retina (211.13±24.37 per mm^2^). To investigate the proportion of spgRGCs in all SC projecting RGCs, we injected AAV2/2-Ef1α-EYFP into the SC site of C57BL/6J mice (n=4). Four weeks later the retina tissues were collected and the labelled cells were counted (Center: 466.79±95.1 per mm^2^; Midperiphery: 960.71±134.04 per mm^2^; Periphery: 1604.51± 302.42 per mm^2^) (Data not shown). The percents of GABA-positive cells in the SC-projecting RGCs were 32.73%, 21.98% and 13.74% in the central, midperipheral and peripheral retina. 74% of EYFP-labeled spgRGCs had the soma diameter smaller than 12.5 μm, which was considered to be a small RGC soma [27]. Based on stratification patterns and light-induced responses, RGCs are classified into ON, OFF, and ON-OFF subtypes. spgRGCs can also be assigned into ON, OFF, and ON-OFF subtypes [28] (ON cells: n=5; OFF cells: n=3; ON-OFF cells: n=4) based on stratification and induced-response pattern (Figs 3G-I). Thus, spgRGCs exhibit preferential periphery distribution in the retina.

**Fig 3.**
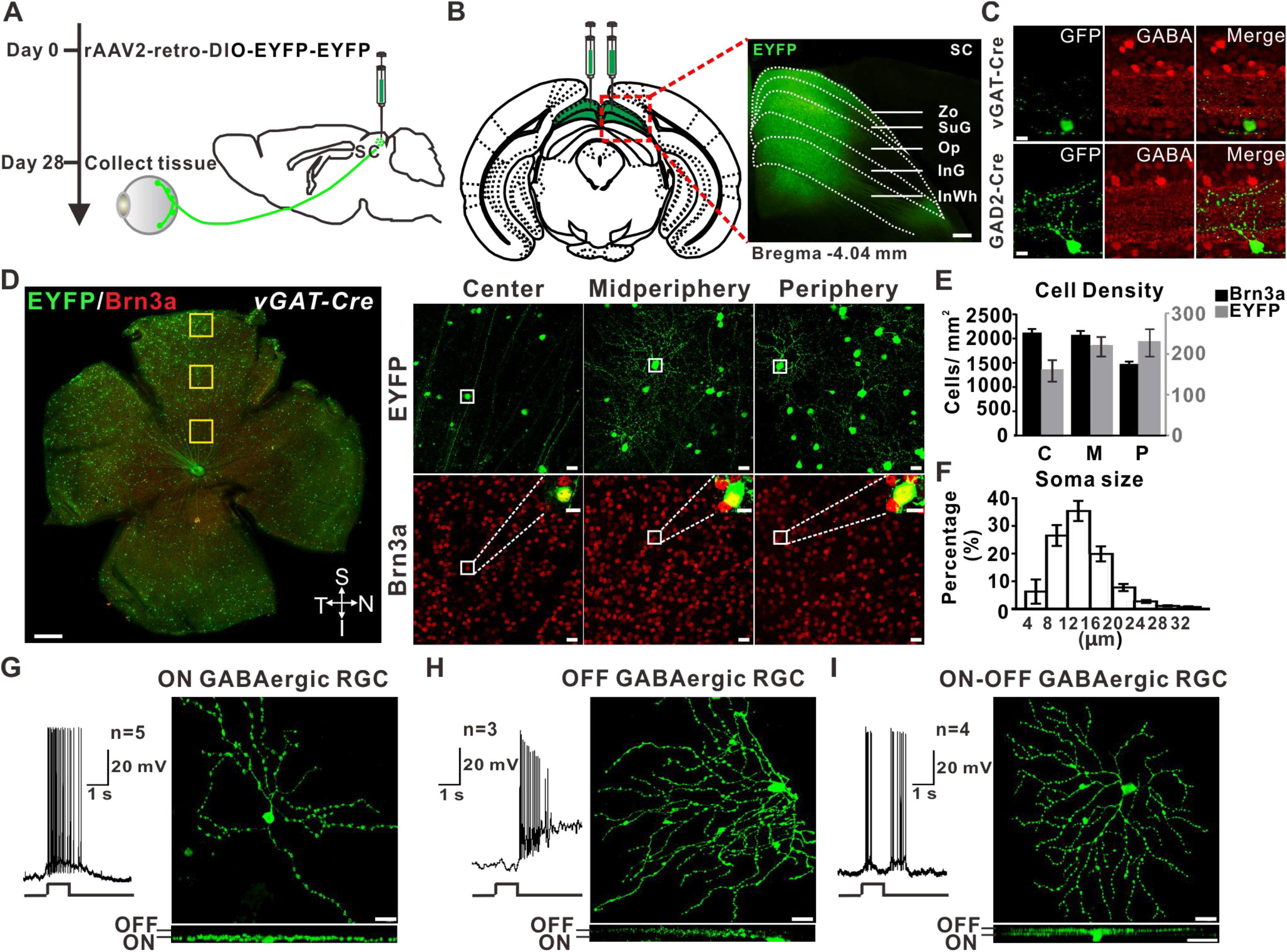
Morphological and functional characterization of spgRGCs. (**A, B**) Experimental design: *AAV-EF1a::DIO-EYFP-EYFP* viruses are delivered into the SC to retrogradely trace spgRGCs. Scale bar: 200 μm. (**C**) EYFP-labeled RGCs also express GABA. Scale bar: 20 μm. (**D**) Representative images of the distribution of EYFP (**upper**) or Brn3a (**lower**) positive RGCs in the flat mount retina of *vGAT-Cre* mice (Scale bar: 500 μm). Highly magnified regions (Scale bar: 20 μm) and counterstained cells (Scale bar: 10 μm) are shown in the right. (**E**) Retinal distributions of the SC-projecting GABAergic RGCs versus Brn3a^+^ RGCs (n=24; means ± SEM). (**F**) Histogram showing the soma size distribution of the SC-projecting GABAergic RGCs. (**G-I**) Light responses and dendritic morphologies of the SC-projecting ON (**G**), OFF (**H**) and ON-OFF (**I**) GABAergic RGCs. Top: flat view; bottom: side view. Scale bar: 20 μm.

### spgRGCs mediate the looming-evoked flight response

To further determine the downstream neurons of spgRGC-projecting SC neurons, we injected the herpes simplex virus (HSV) expressing double-floxed four copies of tandemly repeated GFP genes (HSV-LSL-TK-G4) into the vitreous of *vGAT-Cre* mice to label trans-synaptic neurons (Fig 4A). HSV-LSL-TK-G4 as an anterograde transsynaptic virus tool, is capable of identifying potential postsynaptic targets and facilitates the identification of the cell types. GFP was detected in the SC, the PBG nucleus, the DLG (the relay center in thalamus for the visual pathway) and the pontine nucleus (Pn; involved in motor activity) in the brain (Fig 4B). In the SC, the trans-synaptically labeled SC neurons were parvalbumin-positive (PV^+^) (Fig 4B). These results indicate that spgRGCs are synaptically connected to PV^+^ SC neurons, which are known to form the circuitry controlling the looming-induced flight behavior [4, 9], suggesting that spgRGCs might be involved in the looming-evoked flight response.

**Fig 4.**
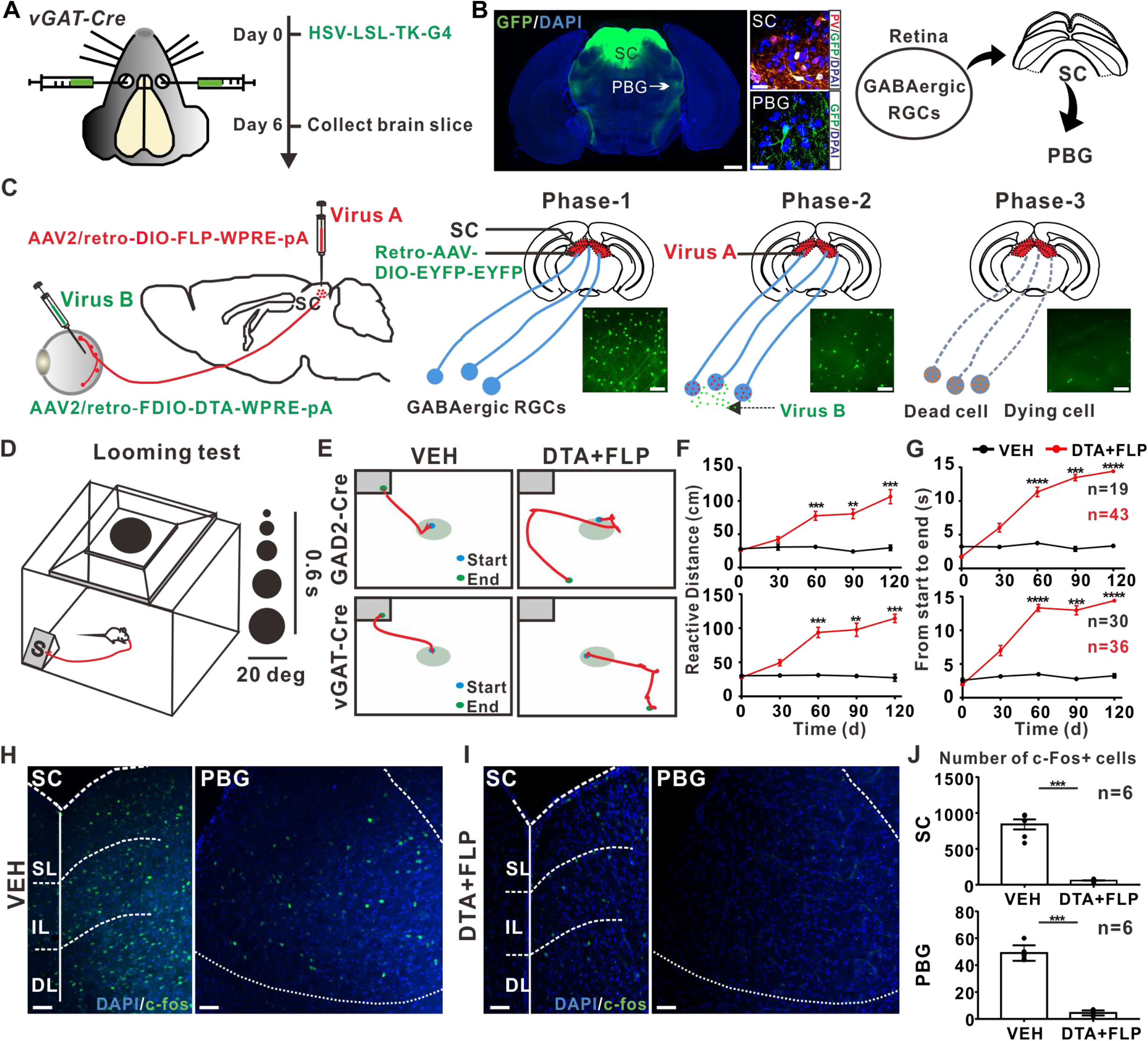
spgRGCs are required for the looming-evoked flight response. (**A**) Experiment design for the trans-synaptic labeling of GABAergic RGCs. (**B**) By infecting the *Gad2-Cre* or *vGAT-Cre* mouse eye with Cre-dependent HSV viruses, the SC, the PBG and the Pn in the brain are labeled by GFP. (Left: Scale bar: 500 μm; Right: Scale bar: 10 μm). (**C**) Experimental strategy for ablating the SC-projecting GABAergic RGCs (phase 1: Expressing FLP in the EYFP-labeled GABAergic RGCs; phase 2: DTA expression via the FLP/FDIO system; phase 3: DTA-induced apoptosis). The SC-projecting GABAergic RGCs can be effectively eliminated. Scale bar: 50 μm (**D-G**). With the expanding looming stimulus at 2-20 degrees for 0.6 second, representative movement traces of the control (VEH) and DTA-expressed (DTA+FLP) *vGAT-Cre* (top, **E**) and *Gad2-Cre* (bottom, **E**) mice. **F** and **G**: quantification results on the movement distance and the time interval from the start to the end point, respectively. (**H-J**) c-Fos positive cells in SC and PBG in looming stimulated control mice (**H**) and DTA+FLP treated mice (**I**). **J**: quantification results on c-Fos positive cells in SC (top) and PBG (bottom) in each group of mice. Scale bar: 50 μm. Mann-Whitney U test; **: *P*<0.01, ***: *P*<0.001, ****: *P*<0.0001.

To directly test their requirement for the looming-evoked flight behavior, we sought to selectively ablate spgRGCs by expressing diphtheria toxin subunit A (DTA), which inactivates elongation factor 2 to terminate protein synthesis and thus kill DTA-expressing cells [25, 29]. AAV2/2-fDIO-DTA was injected vitreously and AAV2/retro-Ef1α-DIO-FLP was injected into the SC of the *vGAT-Cre* or *Gad2-Cre* mice, to simultaneously infect the EYFP-labeled spgRGCs for allowing the Cre-dependent expression of FLP that then drives FLP-dependent DTA expression specifically in spgRGCs(Fig 4C). All injected animals were preselected for fast return to shelter through two rounds of looming stimulus. Two months after dual virus infection, EYFP-labeled spgRGCs have been reduced by approximately 90% (before ablation/phase 1: 204±10.68 EYFP-positive spgRGCs/mm^2^, n=11; after ablation/phase 3: 25.97±2.17 EYFP-positive spgRGCs/mm^2^, n=10; *P*<0.001). In the looming stimulus-induced flight behavior test, there was a significant increase in both “the time to the shelter” and “the total distance” for the mice depleting spgRGCs (Figs 4D-G, Supplemental Movie 1-4). Moreover, the defective looming-evoked flight response caused by spgRGC ablation persists 4 months after the ablation (Figs 4F and 4G). Consistently, the ablation of spgRGCs significantly decreased the number of looming stimulation-induced c-Fos-positive neurons in both the SC and the PBGN, further supporting that spgRGCs are required for looming-induced neuronal activities in the SC-PBGN pathway (Figs 4H and 4I). c-Fos expression is indicative of neuronal activation upon looming signaling-induced stimulation. Our findings indicate that spgRGCs are required for the looming-evoked flight response by activating the SC-PBGN pathway.

### spgRGCs are dispensable for image forming functions

Electroretinography (ERG) and the optomotor response were used to investigate if spgRGCs are required for image-forming functions. There were no significant changes in the a-wave and b-wave amplitudes of scotopic ERGs between the control and ablation groups, indicating that spgRGCs are dispensable for overall retinal functions (Figs 5A and 5B). Although the a wave and b wave correspond to photoreceptor and bipolar cell activity, and therefore should not be impaired by a lack of spgRGCs. The optomotor response was used to determine visual acuity based on head movements in response to the moving vertical grating stimulus. There was no significant difference in the visual acuity between the control and ablation groups (Figs 5C-G). These results indicate that spgRGCs are not required for visual image formation.

**Fig 5.**
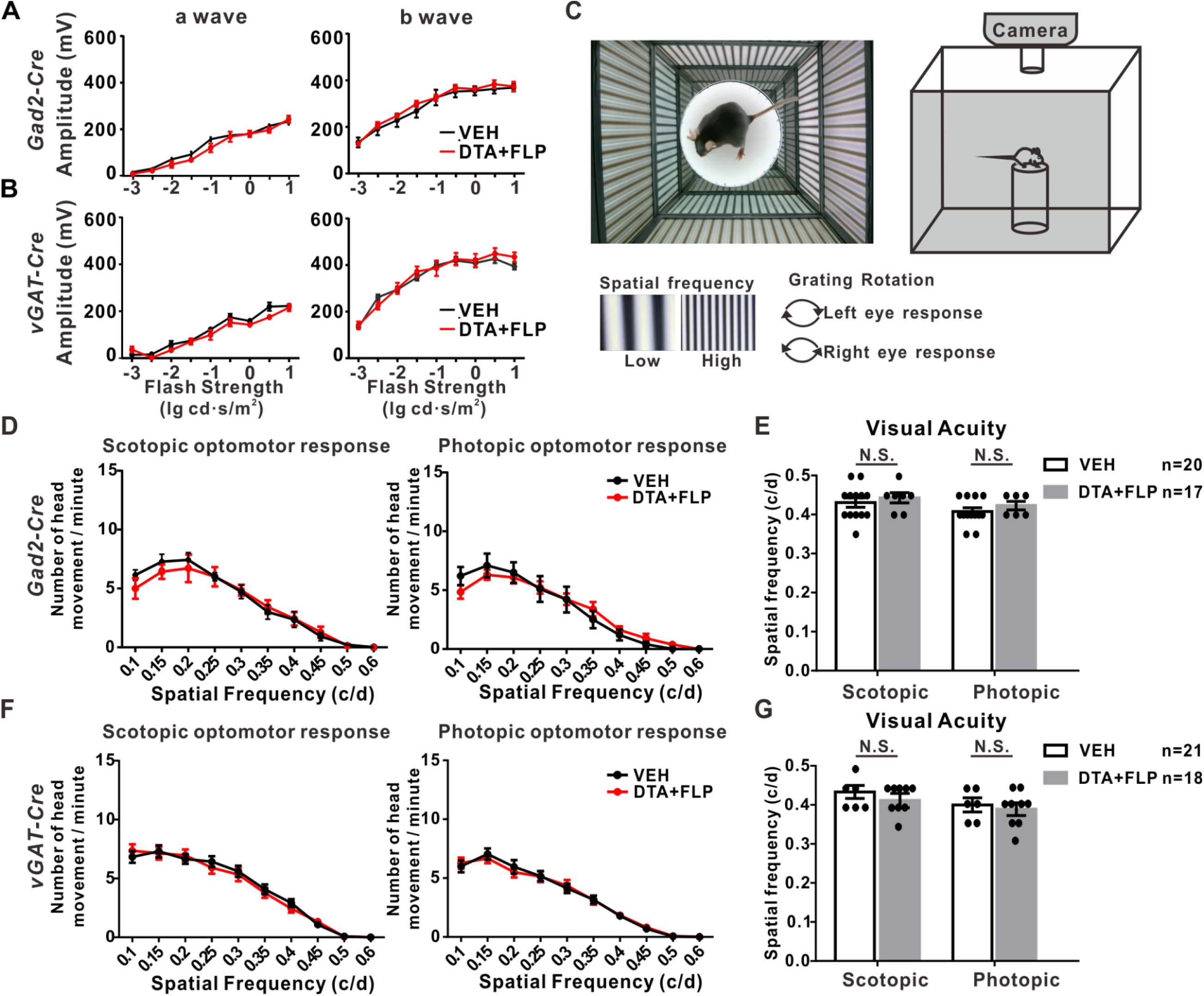
spgRGCs are dispensable for innate visual functions. (**A, B**) ERG amplitudes of a-wave (left) and b-wave (right) show no significant changes in the ablation groups. (**C-G**) A grating stimulation-based OMR setup (**C**), there are no significant differences on the optomotor response (**D, F**) and the maximum spatial frequency (visual acuity; **E, G**) under scotopic and photopic conditions in the control or DTA+FLP induced ablation groups.

### Inhibition of spgRGCs suppresses the looming-evoked flight response

To determine whether inhibitions of SC-projecting GABAergic RGCs are required for looming-evoked response, using the dual-virus intersectional strategy we chemogenetically inhibited spgRGCs by injecting rAAV-DIO-FLP which encoding Cre-dependent FLP bilaterally into the SC and at the same time rAAV-FDIO-hM4D-EGFP (n=5) which encoding FLP- dependent hM4D-EGFP, or rAAV-FDIO-EYFP (n=3) as the control (VEH), into the vitreous chamber bilaterally of vGAT-cre mouse, which upon FLP excision would result in expression of hM4D in spgRGCs (Fig 6A). All injected animals were preselected. Post hoc histological analysis revealed that the hM4D-EGFP expressed in the whole retina (Fig 6B). Behaviorally, we found that CNO administration significantly increased the total duration in hM4D mice but not in the vehicles, furthermore the looming-evoked response of partial animals vanished due to they did not return to shelter during entire stimulation 14.6 s. Taken together, these results demonstrate that inhibition of spgRGCs suppresses looming-evoked flight response.

**Fig 6.**
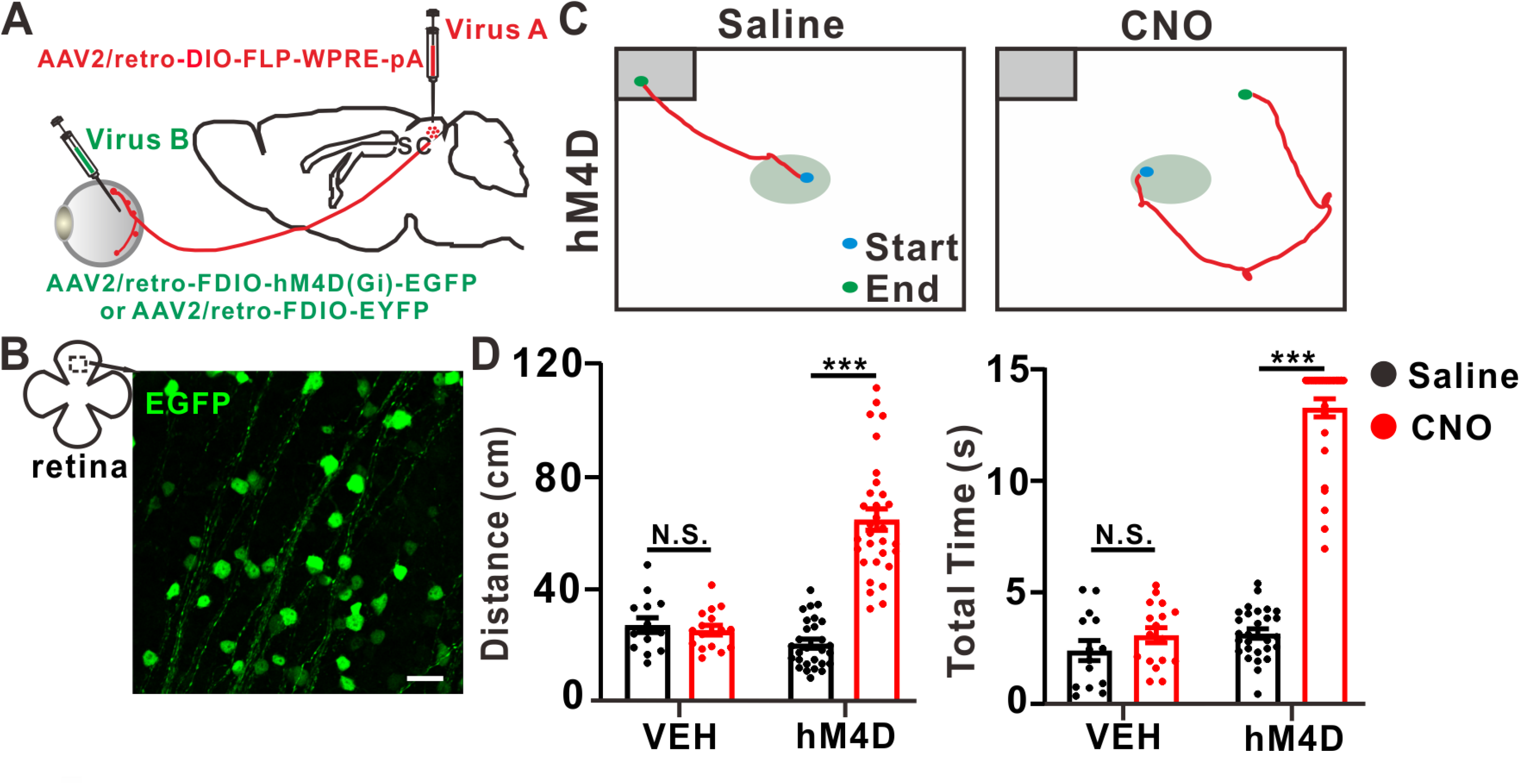
Chemogenetic inhibition of spgRGCs suppresses looming-evoked response. (**A**) Schematics of the dual-virus intersectional approach to target spgRGCs. (**B**) Representative images showing hM4D expression as revealed by immunostaining of GFP in the flat mount retina. 20X, Scale bar: 30 μm. (**C**) Typical movement traces of the saline and CNO groups. (**D**) Quantification results on the movement distance and the time interval from the start to the end point of the vehicle and hM4D groups. Mann-Whitney U test; ***: *P*<0.001.

Aiming at removing the effect of other neurotransmitters except GABA of spgRGCs, AAV with short hairpin (shRNA) was adopted to knock down vGAT expression by injecting rAAV-U6-Loxp-CMV-EGFP-SV40-polyA-Loxp-shRNA (vGAT shRNA group) into the vitreous of the vGAT-Cre mice (S2A Fig). In order to assess the effectiveness and tissue-specificity of the vGAT-Cre combining with LoxP-shRNA system, western blot analysis indicated that the system was in position to efficiently silence vGAT in retina. Retina transfected with vGAT shRNA in two months manifested an inferior expression of vGAT protein to the Control (scramble shRNA group) (S2B Fig). Afterwards, the looming behavior produced by knockdown vGAT in the GABAergic RGCs was compared, vGAT shRNA group conveyed an increased escape latencies and duration during the trials in contrast with the Control group (S2C-D Figs, Supplemental Movie 5-6). In addition, there was no significant diversity between the Scramble and the vGAT shRNA group in the optomotor response (S2E Fig). Nevertheless, specificly knockdown vGAT in spgRGCs remains to be confirmed.

## Discussion

The ability to detect rapidly approaching objects, which mimic predators, is a highly conserved innate defensive behavior in *Drosophila*, navïe rodents [30] and primates [31]. Although the looming-evoked neural circuits in the mouse brain have been nicely studied, it remains poorly understood how the visual looming stimulus is transmitted from the retina to the brain. Here, we show that mouse spgRGCs directly innervate PV^+^ SC neurons to mediate the looming-evoked flight behavior through the SC-PBGN pathway. In the adult mouse retina, GABAergic RGCs project their axons to multiple brain areas, including SC, SCN, OPN and LGN. Moreover, approximately 13% RGCs are GABAergic and belong to ON, OFF and ON-OFF RGC subtypes. In addition, the SC-projecting GABAergic RGCs can release the inhibitory GABA to their mono-synapsed SC neurons when activated. Finally, DTA-mediated ablation of those spgRGCs causes the loss of the looming-evoked flight response. Therefore, we identify a novel GABAergic RGC population in the mouse retina that projects and releases GABA to the SC, and have further showed that spgRGCs function as a part of the neural circuitry mediating the looming-evoked defensive behavior.

In the mammalian eye, RGCs are the only cell type sending long axons to connect the retina to the brain. Earlier studies suggest the existence of GABA-like immunoreactive RGCs in turtle [13], bufo [32] and rabbit retinas [14]. Since GABAergic neurons are believed to function within short distances to regulate local neural networks [33–35], lately a subset of ipRGCs were showed to release GABA and affected non–image-forming behaviors including pupillary light reflex and circadian photoentrainment [12]. In this study, we have provided three pieces of experimental evidence to demonstrate the existence of GABAergic RGCs in the adult mouse retina. First, *Gad2-Cre* and *vGAT-Cre*-driven Cre-dependent expression of EYFP reliably label RGCs that project axons to the brain. *Gad2* and *vGAT* (also known as *Slc32a1*) encode a GABA synthesizing enzyme and a vesicular GABA transporter, respectively, which are both known to be expressed in GABAergic neurons [17, 19, 36]. Consistently, our EYFP-labeled RGCs also GABA positive. Second, we have found that 13% of RGCs express *Gad2* by re-analyzing the recently published RGC SC-seq dataset [37]. Interestingly, most of the 40 RGC subgroups identified by SC-seq, including melanopsin-positive intrinsically photosensitive RGCs (ipRGCs), also express *Gad2* mRNA. Third, optogenetic activation drives EFYP-labeled RGCs to release GABA and elicit the IPSC responses in the projected SC neurons, indicating that those EYFP-labeled RGCs are bona fide GABAergic neurons. In addition, EYFP-labeled GABAergic RGCs project their axons to brain regions as SC, LGN, SCN and OPN. The SC is required for both non-image-forming and image-forming functions, whereas the LGN is required for only visual image-forming functions [38, 39]. The SCN and OPN brain areas are required for non-image-forming functions, circadian photoentrainment [40–42], and the pupillary light reflex (PLR) [12, 43], respectively. The LGN is required for visual image-forming functions, whereas the SC is required for both non-image-forming and image-forming functions [38, 39]. Taken together, multiple types of GABAergic RGCs exist in the adult mouse retina and likely have distinct biological functions.

This study functionally demonstrates that spgRGCs participate in the looming-evoked innate flight response via the SC-PBGN pathway. Based on stratification within the IPL and light-mediated responses, spgRGCs are composed of ON, OFF and ON-OFF types. Those spgRGCs primarily target the superficial and intermediate layers of the SC, which are known to be required for the looming-evoked defensive behavior [8, 44]. Indeed, DTA-mediated ablation of spgRGCs diminishes the looming-evoked flight response, but does not affect image-forming and other non-image-forming functions. Dual SC/DRN-projecting RGCs have been previously identified for the looming-evoked flight response[45]. Since newly identified spgRGCs do not project to the DRN, SC/DRN-projecting RGCs and spgRGCs represent two distinct RGC populations for mediating the looming-evoked flight response. Our trans-synaptic tracing results indicate that the GABAergic RGC-projecting SC neurons are also PV^+^ and are capable of innervating PBGN, PN and DLG in the brain, which should be the same as the previously identified PV^+^ glutamatergic SC neurons [4]. Those PV^+^ SC neurons activate the PBGN-amygdala circuitry to control the looming-evoked flight response [4, 9]. Results of c-Fos expression also support that spgRGCs activate PV^+^ SC and PGBN neurons to mediate the looming-evoked flight response. Therefore, this study shows that spgRGCs are mono-synaptically connected to the SC circuitry for mediating the looming-evoked flight behavior through GABA.

## Materials and Methods

### Animals

The mouse strains used: C57BL/6, *vGAT-Cre* [*B6J.129S6(FVB)-Slc32a1*^*tm2(cre)Lowl*^/ *MwarJ*; Jackson Laboratory] and *Gad2-Cre* (*B6J.Cg-Gad2*^*tm2(cre)Zjh*^/*MwarJ*; Jackson Laboratory). All mice were group-housed and maintained on a 12h light/12h dark cycle with food and water provided *ad libitum*. 6-8-week old adult males were randomly divided into experimental and control groups. All the experiments were done in accordance with the guidelines of Wuhan University Laboratory Animal Care Committee.

### Stereotaxic injection of AAV and HSV

For cell-type-specific retrograde tracing, animals were first anesthetized by intraperitoneally injecting ketamine and xylazine (60 and 16 mg/kg body weight, respectively). Following the making of a small craniotomy hole by a dental drill (OmniDrill35, WPI), a micropipette controlled by a Quintessential Stereotaxic Injector system (Stoelting, Wood Dale, IL, USA, Micro4; WPI, Sarasota, USA) was used to inject 700 nl *AAV2/retro-EF1a-DIO-EYFP-EYFP-WPRE-pA* (6.48 × 10^12^ vg/ml, BrainVTA, Wuhan) virus into the SC of control, *Gad2-Cre* and *vGAT-Cre* mice. Coordinates targeting the SC were as follows: 3.9 mm back from the bregma (anteroposterior), 0.45 mm lateral to the midline (mediolateral), and 1.7 mm deep (dorsoventral) below the pial surface. The pipette was held in place for 10 min, and then withdrawn slowly [46].

For anterograde RGCs tracing, a small incision was made in the conjunctiva to expose the sclera, and 1.5 μl of *AAV2/2-EF1a-DIO-EYFP-EYFP-WPRE-pA* (2.12 × 10^12^ vg/ml, BrainVTA, Wuhan) suspension was injected into the center of the vitreous cavity through the ora serrata with a Hamilton syringe (Hamilton Company, Reno, NV). For anterograde trans-synaptic tracing, 1.5 μl of *HSV-LSL-TK-G4* (1 × 10^10^ vg/ml) suspension was injected into the vitreous chamber, and brain slice were obtained at 6 days [47]. For ablating SC-projecting GABAergic RGCs, two viruses were performed sequentially. 700 nl of *AAV2/retro-EF1a-DIO-FLP-WPRE-pA* (7.33 × 10^12^ vg/ml, BrainVTA, Wuhan) was first injected into the SC of *vGAT-Cre/Gad2-Cre* mice, and 1.5 μl of *AAV2/2-EF1α-FDIO-DTA-WPRE-pA* (6.2 × 10^12^ vg/ml, BrainVTA, Wuhan) or PBS (as a control) was then injected into the vitreous chamber bilaterally 3 days later. GABAergic RGC degeneration were detected after one month [46, 48]. Viral-mediated transfection of DREADD receptors. A chemogenetic approach was used to specifically inhibit spgRGCs. 700 nl of *AAV2/retro-EF1a-DIO-FLP-WPRE-pA* (7.33 × 10^12^ vg/ml, BrainVTA, Wuhan) was first injected into the SC, and 2 μl of *AAV2/retro-nEF1α-FDIO-hM4D(Gi)-EGFP-WPRE* (5.31 × 10^12^ vg/ml, BrainVTA, Wuhan) or *AAV2/retro-nEF1a-FDIO-EYFP-WPRE-pA* (5.35 × 10^12^ vg/ml, BrainVTA, Wuhan; as a control) was then injected into the vitreous chamber 3 days later [49]. All the experiments with viruses were performed in bio-safety level 2 (BSL-2) laboratory and animal facilities.

To remove the putative GABAergic regulation on looming-evoked flight response from spgRGCs, we used vGAT-selective hort hairpin (shRNA) constructs to knock down vGAT expression in the retina by injecting vectors into the vitreous of the *vGAT-Cre* mice. The modified U6 promoter harbors a LoxP-CMV-EGFP-LoxP inside. When the vGAT positive cell was coinfected with LoxP-shRNA vectors, Cre-mediated recombination results in subsequently cuts the LoxP-CMV-eGFP-LoxP inside the U6 promoter and activates the U6 promoter. The activated U6 promoter then drives the down-stream shRNA expression and silences the target gene. rAAV-U6-Loxp-CMV-EGFP-SV40-polyA-Loxp-shRNA (shvgat) and rAAV-U6-Loxp-CMV-EGFP-Loxp-shRNA (scramble/control) were injected into the vitreous chamber. AAVs carrying an shRNA outside two loxP sites (AAV-U6-LoxP-CMV-GFP-LoxP-shRNA, 6.3 × 10^12^ vg/ml) (BrainVTA, Wuhan) were used for Cre-dependent silencing of vGAT in the *vGAT-Cre* mice. The shRNA oligonucleotides for targeting vGAT mRNA (shvgat; 5′-TGCTGTTG ACAGTGAGCGCGGTGTGCTCGTGGTGAATAAGTAGTGAAGCCACAGATGTACTTATTCACCACGAGCACACCATGCCTACTGCCTCGGA-3′) and control shRNA (Control; 5′-TGCTGTTGACAGTGAGCGGCCGCGATTAGGCTGTTATAATAGTGAAGCCACAGATGTATTATAACAGCCTAATCGCGGCTGCCTACTGCCTCGGA-3′) were created following the information described in the literature [50–52]. Control shRNA sequences were used to construct a non-targeting control virus. Mice were given injections of either vGAT shRNA or non-targeting scramble shRNA (control shRNA).

### Histological procedures

For retina whole mounts, eyes were enucleated after cervical dislocation, and eye cups were fixed in 4% paraformaldehyde (PFA; pH 7.4) at room temperature for 40 min [53]. Retina was isolated from the eye cup and flattened. For retina cryosections, eyecups were fixed in 4% paraformaldehyde for 40 min, washed with PBS three times for 5 minutes, incubated sequentially for 1 h in 10% and 20% sucrose, overnight in 30% sucrose. Then the eyes were embedded in O.C.T compound (Sakura Finetek, Japan) and frozen at −80°C. Retinal sections were cut at 14 μm thickness on a freezing microtome (Leica, CM1950, Germany) and mounted on gelatin-coated slides [54]. For brain cryosections, after transcranial perfusion with 0.9% saline followed by 4% paraformaldehyde in 0.1 M of PBS, the brain was removed and post-fixed with 4% paraformaldehyde for 1 day at 4 °C, then transferred into a 30% sucrose solution for 3 days until sectioning with a cryostat. A series of 50 μm sections were collected for staining.

### Immunostaining

The procedures of immunohistochemistry refer to the previous work [55] (the retina and brain slices were first incubated for 2 hours in a blocking solution containing 5% bovine serum albumin (BSA), and 0.2% Triton X-100 in PBS (pH 7.4). And then incubated in primary antibodies at 4°C overnight. After washing with PBS for 3×15 min, the samples were incubated in the appropriate secondary antibodies for 2 hours at room temperature. The samples were washed again in PBS for 3×15 min and mounted in DAPI (4-,6-diamidino-2-phenylindole; Life Technologies, D1306) for 5 min. The antibodies used in this work were list as follows: chicken polyclonal antibody against GFP for immunolabeling EYFP^+^/GFP^+^ neurons (1:2000, Abcam, ab13970); mouse polyclonal antibody against Brn3a (1:500, Millipore, MAB1585); goat polyclonal antibody against ChAT (1:200, Millipore, AB144P); rabbit polyclonal antibody against GABA (1:200, Sigma, A2052), rabbit polyclonal antibody against c-Fos (1:500; PC38T, Calbiochem). Secondary antibodies were anti-isotypic Alexa Fluor conjugates (1:500, Invitrogen), incubation for 2 h at room temperature, and after washes in PBS, the retinas were subsequently mounted in antifade mounting medium.

### Confocal microscopy and three-dimensional reconstruction

Images were collected with a confocal laser-scanning microscope (Leica, SP8) and Brightfield, Fluorescence & FISH Digital Pathology Scanner (Leica, Aperio VERSA 8). For three-dimensional reconstruction of injected or virus labelled RGCs, the Z-axis interval was set at 0.4 μm and areas of interest were scanned with a 100X oil immersion objective. A montage of optical section stacks was created, projected to a 0°X-Y plane and a 90°X-Z plane to obtain a 3-D reconstruction. Each stack of optical sections covered a retinal area of 325.75 × 325.75 μm^2^ (1,024×1,024 pixels). Details of three-dimensional reconstruction and confocal calibration procedures were described elsewhere [56]. Contrast and brightness were adjusted. Total soma and dendritic field size of each filled cell were analyzed.

### Electrophysiological recording on tissue slices

Brain slices containing the SC were prepared recording to the procedures described previously [45]. In brief, mice were anesthetized with isoflurane. After decapitation, brains were immediately immersed in ice-cold oxygenated (95% O_2_ and 5% CO_2_) cutting solution which containing (in mM): Choline chloride 97, Ascorbic acid 11, KCl 2.5, NaHCO_3_ 25, NaH_2_PO_4_ 1.26, Sodium Pyruvate 3, CaCl_2_ 0.5, MgCl_2_ 7.2, glucose 25. Brain slices (350 μm) was cut using a vibratome (VT1200S, Leica, Germany). The slices were incubated in 34°C artificial cerebrospinal fluid (ACSF: NaCl 118, KCl 2.5, CaCl_2_ 2, NaH_2_PO_4_ 1, MgCl_2_ 2, NaHCO_3_ 26 and glucose 22 in mM) and bubbled with 95% O_2_ and 5% CO_2_ for 30 minutes before transfer to the recoding chamber. After an hour incubation at room temperature (25°C), slice was transferred to the recording chamber, visualized with an X40 water immersion lens, DIC optics (BX51WI, Olympus, Japan) and a CCD camera (IR1000, DAGE-MTI, USA). The data were digitized and recorded on-line using Clampex 10.6 (Axon Instruments, Molecular Devices). Signals were amplified by MultiClamp 700B (Molecular Devices), filtered at 1 kHz and sampled at 10 kHz. Electrodes were fabricated from borosilicate glass (Sutter instrument) using a pipette puller (Sutter, P1000); with resistance 5-7 MΩ. The internal solution for the recording glass pipette was (in mM): Cs-methanesulfonate 115, CsCl 20, HEPES 10, MgCl_2_ 2, ATP-Mg 4, GTP-Na 0.4, Na-phosphocreatine 10, EGTA 0.6 (pH 7.2 adjusted by CsOH, mOsm 290). After establishing the whole-cell configuration, the holding potential was maintained at 0 mV and GABA receptor-mediated IPSCs were pharmacologically isolated using an antagonist cocktail (D-AP5 50 μM; strychnine 10 μM, CNQX 10 μM and TTX 1 μM) [57]. ChR2-induced synaptic current was recorded while holding at −70 mV. The external solution contained NaCl 125, KCl 2.5, CaCl_2_ 2, MgSO_4_ 1, NaHCO_3_ 26, NaH_2_PO_4_ 1.25 and Glucose 20. All drugs were obtained from Sigma-Aldrich or Tocris Bioscience. For light stimulation, an optic fiber was positioned right above the slice (Mightex, BioLED Light). Synchronized light-evoked response was triggered by Clampex 10 to deliver light pulses (470 nm, 5 ms, 0.05 Hz). For retinal slice recording, the isolated retinas were cut into 15-μm-thick slices in Ringer’s using a manual cutter (ST-20, Narishige, Japan). The slices were transferred into a recording chamber with the cut side up and held in place with vaseline. Whole-cell recordings were obtained from selected neurons. ChR2 positive GABAergic neurons was confirmed by enhanced yellow fluorescent protein (eYFP) labeling.

### CNO administration

The DREADDs program involves injecting an engineered G protein-coupled muscarinic receptor modified to respond only to the synthetic compound clozapine N-oxide (CNO) which activates the DREADD producing the spgRGCs inhibition [58]. CNO (Absin, abs812140) was dissolved in DMSO then diluted to a final concentration of 5 mg/ml CNO in 0.5% DMSO in saline solution. Vehicle injections were 0.5% DMSO in saline solution [59]. To confirm the specificity of DREADD-CNO interactions, DREADD-free mice with EYFP-CNO group was included. Animals received either equal volume vehicle or CNO (5 mg/kg, i.p.) 30 min before the behavior assays [49].

### Looming stimulation test

The looming stimulation test was performed in an open-top acrylic box, as described by Huang L, et al [6]. The arena had dim lighting from the screen of the monitor. The floor and three walls were covered with a frosted coating to prevent reflections of the stimulus. A camera from the side was used to track the movements. An LED monitor was embedded in the ceiling to present the looming stimulus. The stimulus program was displayed on a LED monitor (HPZ24i, HP, USA). The looming stimulus, which consisted of an expanding black disc, appeared at a diameter of a visual angle from 2° to 20° in 0.6 s, for 15 times in 14.6 s. The stimulation was initiated manually when the mouse was in the center of the arena [5, 6, 48]. Looming tests were conducted in Control mice (n=22 for *Gad2-Cre* mice; 28 for *vGAT-Cre* mice) and DTA+FLP mice (n=43 for *Gad2-Cre* mice; 35 for *vGAT-Cre* mice).

### Optomotor response

Optomotor responses of animals in Control (n=20 for *Gad2-Cre* mice; n=21 for *vGAT-Cre* mice) and DTA+FLP (n=17 for *GAD-Cre* mice; n=18 for *vGAT-Cre* mice) groups were measured as described [60]. Briefly, mice were placed on a platform (9 cm diameter, 17.5 cm above the bottom of the mirror) surrounded by four LCD screens (Lenovo, L1900 PA) that displayed rotating black-and-white stripes at various spatial frequencies. The platform was set in the middle of the arena for mouse to stand and move freely. Stimuli were presented on four computer screens surrounding the animal covering the whole field of view with a texture projected onto the surface of a virtual sphere. Mice were videotaped with a camera mounted on top of the platform for subsequent scoring of head tracking movements. Animals were dark-adapted for 3-4 hours, and then presented with vertical black and white stripes of a defined spatial frequency. These stripes were rotated alternately clockwise and anticlockwise for 30 s in each direction and paused for 10 s. The animals were tested with spatial frequencies increasing from 0.1 to 0.6 cycles per degree (cyc/deg): 0.1, 0.15, 0.2, 0.25, 0.3, 0.35, 0.4, 0.45, 0.5 and 0.6 cyc/deg. Procedures for measuring optomotor responses under photopic condition were similar to the scotopic condition except that animals were subjected to 400 Lux light stimulus for 10 mins to allow them to adapt to the light.

### Electroretinography

Retinal functions of control (n=16 for *Gad2-Cre* and *vGAT-Cre*) and DTA+FLP (n=24 for *Gad2-Cre* and *vGAT-Cre*) mice were measured with a commercial ERG system (RetiMINER, China). After overnight dark adaptation [61], the mice were anesthetized by intraperitoneally injecting ketamine and xylazine (60 mg/kg and 16 mg/kg bodyweight, respectively), and then lightly secured to a stage with fastener strips across the back to ensure a stable, reproducible position for ERG recordings. All procedures were performed under dim red light. Customized gold-loop wire electrodes were placed on the center of each eye to measure the electrical response of the eye to flash stimulation. Reference electrodes were placed subcutaneously between the ears and a ground electrode was placed in the base of the tail. The ERG consisted of a 9-step series of full-field flash stimuli dark-adapted (−3 to 1 lg cd·s/m^2^) conditions. After the ERG recording, erythromycin was used on the ocular surface, and the animals were allowed to recover on a heating pad [62].

### Western blotting

Western blotting was performed according to previous work on retinal extracts [54]. They were homogenized with RIPA lysis buffer (Thermo Scientific, USA) supplemented with a PMSF (Servicebio, Wuhan). Protein concentration in the supernatant was measured using BCA Protein Assay kit (Servicebio, Wuhan). Proteins (40 μg) in retinal samples were resolved by 10% sodium dodecyl sulphate polyacrylamide gel electrophoresis (SDS-PAGE) and transferred onto and electro-blotted to nitro-cellulose membranes (Millipore, USA). The membranes were subsequently blocked with 5% nonfat milk for one hour and incubated overnight at 4 °C with primary antibodies, including rabbit anti-SLC32A1 (1:1000; Absin, abs136723), rabbit anti-GAPDH (1:6000; Antgene, ANT012). The membranes were washed for 30 min in TBS-Tween 20 and incubated with HRP-conjugated goat anti-rabbit IgG (1:5000, Servicebio) for 2 h at RT. After the second antibody, the membranes were washed again and immunoreactive bands were subsequently detected using ECL reagents and recorded by X-ray films.

## Data quantification and statistical Analysis

All experiments were performed with anonymized samples in which the experimenter was unaware of the experimental condition of mice. Statistical analysis was performed using SPSS software (IBM, Armonk, NY, USA). Unpaired *t-test* or Mann-Whitney U test were used. Error bars represent the mean ± standard error of mean (SEM). p<0.05 is considered as significant difference between two samples [54, 63].

## Author Contributions

Conception and design of the experiments: Y. Shen and X. Luo. Performed the experiments: X. Luo, Q. Deng, D. Cai and K. Shen. Analysis and interpretation of data: Y. Shen, T. Xie, X. Luo, D. Cai, K. Shen and H. Li. Wrote the paper: Y. Shen, X. Luo and D. Cai. All authors approved the final version for publication.

## Supplemental Information

**S1 Fig.**
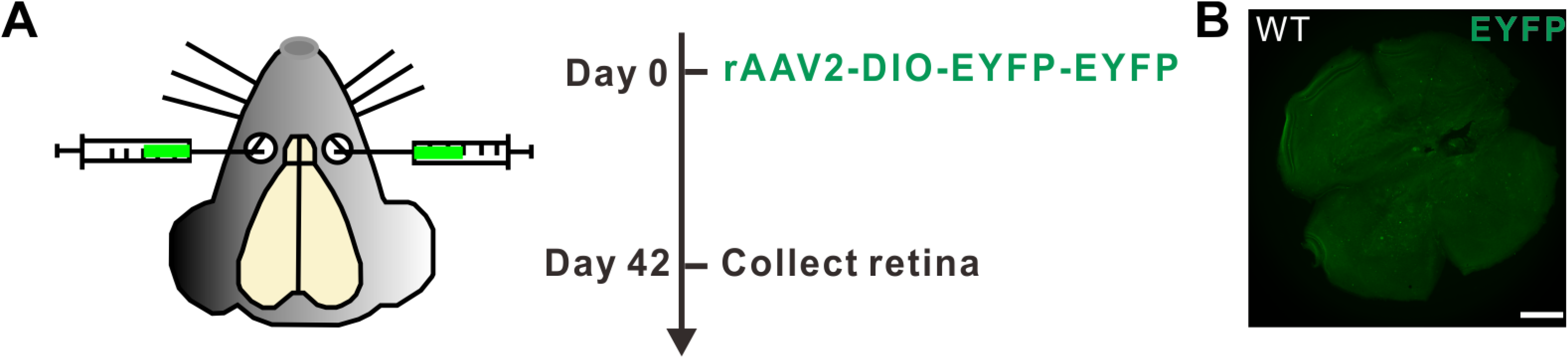
No virus-mediated labeling in control mice following *rAAV2-DIO-EYFP-EYFP* infection. (**A**) Experimental strategy for viral delivery. (**B**) No EYFP-positive cells are detected in the flat-mounted retina 42 days after virus infection, confirming the Cre-depended EYFP expression in this study. Scale bar: 100 μm.

**S2 Fig.**
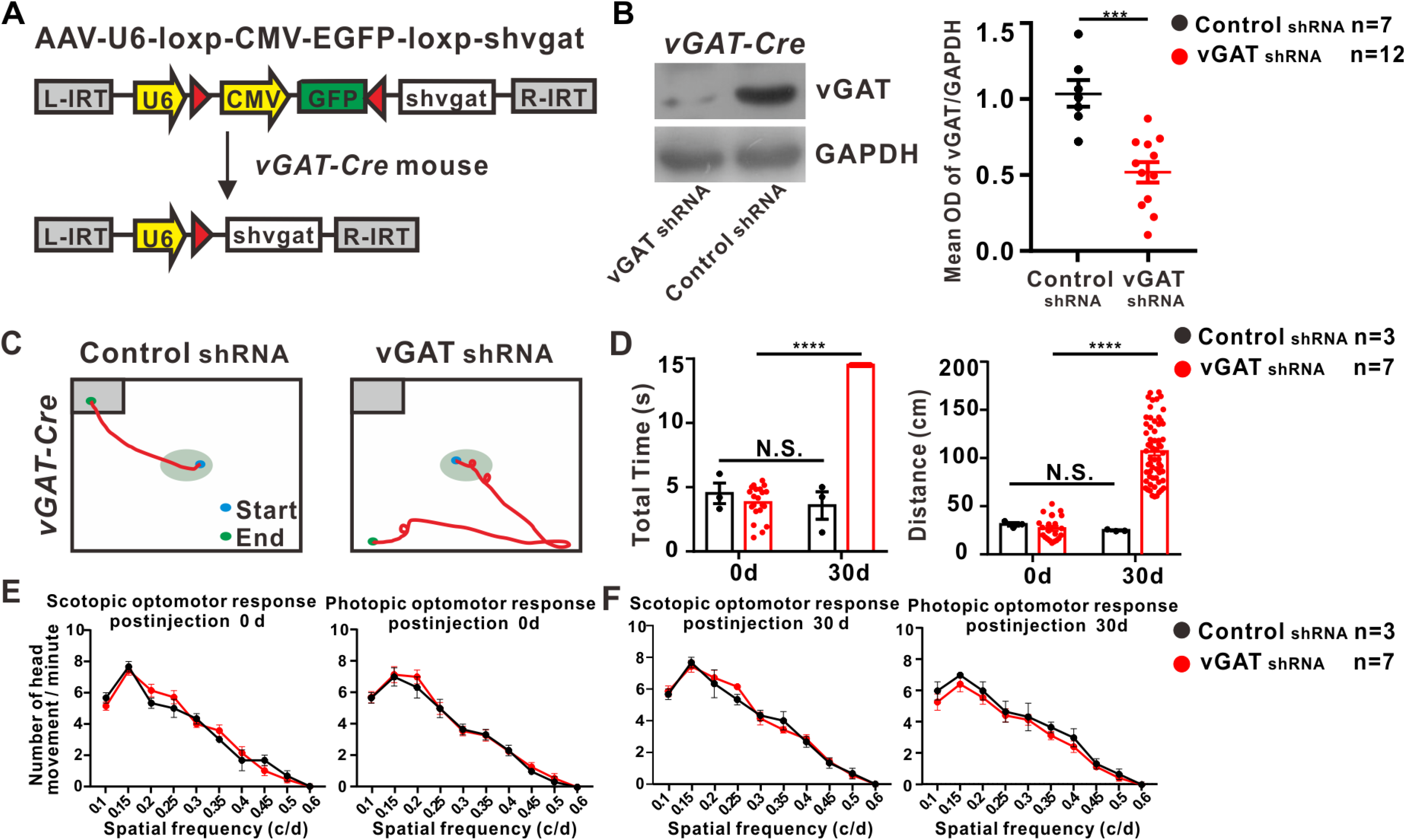
Looming-evoked flight response requires functional release of GABA from RGCs. (**A**) The hairpin (sh) oligonucleotide (sholigo), targeting the vGAT transcript, was used to knock down vGAT expression. (**B**) Western blot analysis show that it can efficiently knockdown vGAT. Representative immunoreactive bands of vGAT protein levels, the bottom panel show the mean optical density (OD) of the band corresponding to vGAT normalized to GAPDH in the two groups. (**C-D**) Typical movement traces of the Control shRNA and vGAT shRNA groups. (**D**) Before or 30 days after injection. Quantification results on the movement distance and the time interval from the start to the end point, respectively. (**E-F**) Functional release of GABA from spgRGCs are dispensable for the optomotor response. Before or 30 days after injection. there are no significant differences on the optomotor response under scotopic and photopic conditions in the Control shRNA or vGAT shRNA group.Mann-Whitney U test; ***: *P*<0.001, ****: *P*<0.0001.

**Supplemental Movie 1**

Example of looming-evoked defensive behavior in *Gad2-Cre* mice.

**Supplemental Movie 2**

Example of looming-evoked defensive behavior in *vGAT-Cre* mice.

**Supplemental Movie 3**

Effects of spgRGCs deletion on looming-evoked flight response in *Gad2-Cre* mice.

**Supplemental Movie 4**

Effects of spgRGCs deletion on looming-evoked flight response in *vGAT-Cre* mice.

**Supplemental Movie 5**

Example of looming-evoked defensive behavior in vGAT shRNA group.

**Supplemental Movie 6**

Example of looming-evoked defensive behavior in control shRNA group.

